# High-throughput SNP discovery, development and validation of a 30 K target SNP genotyping tool for cultivated flax (*Linum usitatissimu*m) breeding and germplasm characterization

**DOI:** 10.1101/2025.09.24.678307

**Authors:** Boris B. Demenou, Christophe P. Pineau, Isabelle Le Clainche, Aurélie Berard, Patricia Faive-Rampant, Damien D. Hinsinger

## Abstract

The cultivated flax (*Linum usitatissimum* L.) is an industrial crop widely cultivated for fiber and seeds, in broad geographical regions around the world. This crop faces many challenges (yield, quality, biotic and abiotic stresses) linked to climate change as almost all other crops. Among solution for improving or breeding new adapted cultivars, marker-assisted selection has been widely applied in plant breeding to enhance crop yield, quality, and tolerance to biotic or abiotic stresses. Recent advance of targeted genotyping-by-sequencing (GBS) offers an ultimate MAS tool to accelerate plant breeding and crop improvement.

To facilitate the utility of SNP-based genotyping, we developed and validated in this study a target SNPs genotyping tool named AT-SNP-30K using Allegro targeted SNPs technology. A total of 41k SNPs were selected from 4.78 million and 3.73 million SNPs identified in two different accessions panels, respectively. Probe design for all these markers was successfully achieved for 35,791 of these markers, representing a 86% conversion success rate. The set of markers was then validated by genotyping a diversity panel comprising 384 individuals, including 376 accessions and eight replicates of the fiber flax Idéo cultivar. The validated genotyping tool includes 35,791 SNPs, covering the fifteen chromosomes with 24,951 high-quality SNPs (MAF> 5%, average low rate of missing data) and 27,247 SNPs having a MAF greater than 1%; demonstrating high polymorphism and excellent genotyping accuracy. The repeatability of genotyping in the validation experiment, reached 99.00% of SNPs for the eight Idéo replicate controls. The AT-SNP-30K genotyping tool is a robust resource for genetic studies, germplasm characterisation and cultivated flax marker-assisted selection studies. It can be used to enhance the breeding of new flax cultivars adapted to the context of climate change.

## INTRODUCTION

Genotyping tools are one of the important tools that can enhance the efficiency of breeding programs through their use in germplasm characterization and in marker-assisted plant breeding. Among genotyping methods, Genotyping-by-Sequencing (GBS, Elshire et al., 2011) is a simple, highly multiplexed approach for constructing reduced-representation libraries for the Illumina Next-Generation Sequencing (NGS) platform (He et al 2014). In recent years, targeted GBS, a GBS technology with a key difference of selecting target markers at an attractive price per sample, has become an essential technology for plant breeding (Endelman et al. 2024). Targeted GBS uses a powerful multiplexed approach that enables users to interrogate hundreds to thousands of markers across hundreds to thousands of samples simultaneously. It includes the design of a set of primer pairs or oligonucleotide baits for targeted markers, the number of which varies depending on the application and price point (Ali et al., 2016; Campbell et al., 2015).

Among the various available molecular markers, the use of single base changes in the sequence, called single nucleotide polymorphisms (SNP) markers, to identify and differentiate between crop cultivars and varieties is widespread (Cull and Joly, 2025), even for GBS technology. SNPs were used at large scale for genotyping new accessions and were made accessible for many reasons: (i) SNP markers are the most abundant source of variation in plant and animal, making them ideal for large-scale genotyping (ii) SNP markers are amenable to high-throughput, cost effective genotyping technologies; (iii) SNP-based marker techniques have been improved in marker density and, if compared with the earlier genotyping approaches, costs and time required for SNP discoveries have been significantly reduced; (iv) SNP markers analysis has the advantages of the high probability of finding a marker within the gene of interest due to its high density across the genome (Syvänen, 2005); (v) for SNP-based genotyping assays, notably that once the initial sequencing has been completed and markers have been selected, simple Polymerase Chain Reaction (PCR) style assays are sufficient to genotype plants; (vi) The development of Next-generation and cost-effective sequencing technologies (Oxford Nanopore and Pacific Biosciences technologies, Illumina and other) favours ultra-high-throughput sequencing and therefore high-quality reference genomes assembly (Hu et al. 2021; Park et al. 2016).

Among the available targeted GBS technologies, the Allegro® Targeted Genotyping V2 technology represents a recently implemented fast, scalable (highly multiplexed experiments) and cost-effective (through a pooling strategy) approach to perform targeted genotyping-by-sequencing on a wide variety of organisms using next generation sequencing (NGS). This approach is based on the Single Primer Elongation Technology (SPET), in which only one primer is required per targeted SNP, reducing locus-drop issues that can be observed in PCR-based approaches when genotyped samples are genetically distant from the reference used for primers design. In agrigenomics, this technology has demonstrated significant promise for various crops (e.g. Barchi et al. 2019; Tripodi et al. 2023; Lippolis et al. 2025) and other cultivated species (e.g. Scaglione et al. 2019; Baccichet et al. 2022). Furthermore, Allegro genotyping provides newly detected SNP close from the target SNP that can improve accessions delineation. Similar to GBS but targeting specific SNPs/loci or gene of interest, the Allegro protocol represents a promising alternative for species that require high-throughput, low-cost genotyping solutions but lack existing SNP arrays (Scaglione et al. 2019). This flexibility makes it especially valuable for orphan crops, non-model species, and emerging agricultural organisms where traditional array-based platforms are not commercially available, enabling researchers to develop custom targeted panels that meet their specific breeding and research objectives while maintaining cost-effectiveness and scalability.

The cultivated flax (*Linum usitatissimum* L.) is an industrial crop widely cultivated in broad geographical regions around the world for fiber and seeds. Fiber flax is one of the top fiber crops used in the textile industry, whereas linseed flax ranked fifth oilseed crop in the world (Ottai et al.2011). Fiber flax is mainly cultivated in Europe (80%) and Asia (20%), while the Americas (41.2%), Asia (35.4%), and Europe (18.5%) were the leading producers of linseed (Soni 2021; FAOSTAT 2022). In Europe, the Benelux countries (France, Belgium and the Netherlands) are the leading fiber flax producer with up to 586 110 t of production (FAOSTAT, 2023). Genetic improvement priorities in flax vary according to its intended use, fiber or oilseed, and are primarily aimed at addressing current agronomic and climatic constraints (Gouy et al, 2025). The breeding efforts are mainly focused on the resistance to major fungal pathogens (*Polyspora lini, Septoria linicola*, fusarium wilt, flax scorch, and powdery mildew), the tolerance to abiotic stresses such as elevated temperatures, drought, but also cold, particularly for winter-type lines, the yield of fiber and seed, the quality of fiber and oil. Breeding improvement will only be possible using, among other, marker-assisted breeding tools. Indeed, many high-quality reference genome assemblies have been published in recent years (see Demenou et al., 2025 for a summary) and have been made available for genetic studies. However, no high-density genotyping tool has been available until now. The most recent flax genetic studies have employed WGS resequencing data from genotype panels (e.g. You and Cloutier, 2020; Kanapin et al., 2021; Speck et al., 2022) or have used markers other than SNPs (e.g. Spielmeyer et al., 1998; You and Cloutier, 2020).

In this study, we report the identification of high-throughput SNP markers in two genotype panels to develop a genotyping tool for flax breeding and genotype characterization. Specifically, we: (i) identified a set of SNPs representative of cultivated flax diversity using two panels of 96 European and 77 worldwide flax genotypes; (ii) selected a set of 40,000 SNP variants and developed a targeted genotyping-by-sequencing tool using the Tecan Allegro targeted Genotyping V2 technology; and (iii) validated this tool by genotyping a panel of 376 flax genotypes and checking its repeatability in eight replicates of the control genotype.

## MATERIAL AND METHODS

### Plant material and panels composition

A first diversity panel comprising 96 flax (*Linum usitatissimum* L.) entries (hereafter panel_96) was established to capture the genetic variability present in contemporary European breeding programs and to include additional sources of worldwide variability. The selection aimed to represent a broad spectrum of phenotypic and genetic diversity, including elite cultivars, lines derived from exotic germplasm, and bridging lines designed to connect distant genetic backgrounds with elite material. The panel_96 includes both fiber flax (spring and winter types) and linseed (spring and winter types), categorized as follows: SFF (spring fiber flax), WFF (winter fiber flax), SOF (spring oil flax), WOF (winter oil flax). The final composition was SFF = 37 (38.5%), SOF = 34 (35.4%), WFF = 8 (8.3%), and WOF = 17 (17.7%). Regarding provenance, entries originated primarily from France (n = 63) and the Netherlands (n = 17), with additional accessions from Romania (n = 2), Canada (n = 1), United Kingdom (n = 1), and 12 entries with unknown country information. By program category, commercial lines (“Line”) accounted for 56 entries, research material (“research_cross”) for 26, and worldwide variability accessions (“TRT”) for 14. Notably, the TRT subset is enriched in SOF types (12 of 14), whereas WFF and WOF are represented exclusively by Line and research_cross categories. Breeding companies represented in the panel include Terre de Lin, Linéa, Laboulet Semences, and Limagrain for France, and Wiersum/Van de Bildt for the Netherlands. The detailed composition of this panel is provided in Supplementary Table S1.

To complete the first panel with worldwide genetic diversity, we selected a second panel of 77 accessions (hereafter panel_77) among the 200 accessions sequenced by Guo et al. (2020). Elite cultivars were excluded as already selected in the first panel. We selected accessions from all countries and geographical regions included in the Guo et al. (2020) collection, as well as from all flax types. Where several genotypes had the same characteristics (e.g. country and type), only one was chosen at random. This panel included 19 fiber flax, 47 oilseed flax and 11 unknown or dual-purpose type, from 46 countries (Supplementary Table S2). Genomic sequences data published by Guo et al. (2020) for the 77 accessions were downloaded from SRA NCBI archive.

To validate the genotyping tool, an extended panel of 376 accessions was selected (hereafter panel_384). It included 178 commercial flax varieties cultivated in Europe over recent decades, chosen for the availability of historical phenotypic data, 198 accessions (Supplementary Table S3) selected among the germplasm collection of 1,650 cultivated flax (fiber, oilseed and dual purposes type) maintained by Arvalis since 2010 and 8 repetitions of Idéo accession to test the repeatability of the genotyping tool. This collection is predominantly composed of spring-type inbred lines, with 66% belonging to the oilseed group, 22% to the fiber group, and 12% classified as dual-purpose (both fiber and oilseed).

### Illumina short read resequencing, *SNP discovery and target SNP selection*

Genomic DNA of all the European flax of the panel_96 was isolated from young seedlings (12–15 days old) using the Mag-Bind® Plant DNA DS 96 Kit (Omega Bio-Tek, ref. M1130-01), following the manufacturer’s recommendations with minor adaptations for high-throughput processing on a TECAN EVO150 liquid-handling platform equipped with a 96-channel head. Plant material was flash-frozen in liquid nitrogen and ground using a Retsch MM200 mixer mill with 3 mm steel beads. DNA quality was assessed spectrophotometrically, and only samples with A260/A280 and A260/A230 ratios between 1.8 and 2.0 were retained. Final DNA concentrations were adjusted to 20 ng·µL^-1^, with a minimum yield of 1000 ng per sample to ensure sufficient input for downstream library preparation. Short reads libraries were prepared from 800-1000ng of genomic DNA following the Kapa Hyper prep PCR free protocol with Illumina TruSeq DNA UD Indexes (Santa Clara, CA, USA). Libraries were quantified by qPCR (MxPro, Agilent Technologies, Santa Clara, CA, USA) with the KAPA Library Quantification Kit for Illumina Libraries (Roche), and their profiles assessed with an Agilent High Sensitivity DNA kit on the Agilent 2100 Bioanalyser. Libraries were then sequenced at EPGV (INRAE, Evry, France) on two lanes of an Illumina NovaSeq6000 – S4 flow cell, with PE 2×150 cycles (Illumina, San Diego, CA, USA). An in-house quality control process was applied to the reads that passed the Illumina quality filters as described in Alberti et al. (2020). Briefly, low-quality nucleotides (Q < 20) from both ends of the reads were discarded, Illumina sequencing adapters and primer sequences removed, and reads shorter than 30 nucleotides after trimming discarded. Finally, read pairs that map to the phage phiX genome were identified and discarded, using SOAP aligner (Li et al., 2008), and the Enterobacteria phage PhiX174 reference sequence (GenBank: NC_001422.1).

The two flax panels panel_96 and panel_77 were used for variant identification, separately. Genomic sequences data for the 77 worldwide panel published by Guo et al. (2020) were downloaded from SRA NCBI. The quality of the raw reads was assessed using FastQC (S.Andrews. 2010), and low quality and short sequences were trimmed using Trimmomatic (Bolger et al. 2014) with the parameters LEADING:20 TRAILING:20 SLIDINGWINDOW:4:30 MINLEN:30.

For variant identification, we used the Idéo assembly as genome reference (Demenou et al. 2025). We mapped the clean reads of the accessions against the Idéo reference using the BWA tool (Li and Durbin, 2009), with default parameters. Prior to variant calling, we used MarkDuplicates (picard tools, https://broadinstitute.github.io/picard) to locate, tag and remove duplicate reads in the BWA BAM output file (where duplicate reads are defined as originating from a single fragment of DNA; duplicates can arise during sample preparation e.g. library construction using PCR). We generated mpileup files for each panel from BAM files using bcftools and samtools (mpileup, Danecek, Bonfield et al. 2021) and then used Varscan v2.4.3 (mpileup2snp, Koboldt at al. 2012) to detect SNP and indel variants with the following conditions: mapping quality score ≥ 20, SNP site depth ≥ 10 (3 reads for panel_77), average base quality score ≥ 20, minor allele frequency ≥ 5 and a p-value of 0.05, default value for the other parameters.

Single nucleotide polymorphism (SNP) variants detected were further filtered from the two variant calling format (VFC) files according to many criteria, with the aim of selecting around 30 to 35K SNP to develop a genotyping tool. First, we retained high-quality SNPs as biallelic SNPS with a missing rate (MR) < 10% (50% for the second panel) and on a minor allele frequency (MAF) ≥ 5%. We then filtered the SNP for (i) read mean depth (minDP=10 and maxDP=100) and mapping quality (--minQ 30 and -- minGQ 30); (ii) heterozygosity (<20%) and genotypic frequency (homozygous genotype with frequency between 5% and 95%); (iii) number of blast hits (only one hit against Idéo genome reference) to ensure the specificity of context sequences and finally for (iv) linkage disequilibrium (only unlinked SNPs with r^2^ < 0.7 were kept).

### Custom probes design for targeted SNP

The selected SNPs were submitted to TECAN for design. The genome of Idéo (Demenou et al. 2025) was used as reference and provided as fasta, with the position of the targeted SNP on this reference submitted as a BED file. The resulting design was assessed for the number of SNP covered by one or two probes, the number of matches of each probe in the reference genome by BLASTN and standard parameters, as well as their distribution on the genome. Once the design validated, the synthesis of *in solution* probes was performed by TECAN.

### SPET Allegro Genotyping tool validation

The genotyping tool was validated by genotyping the panel_384. Genomic DNA of all the accessions of this panel was isolated from young leaves and was quantified as described above.

#### Libraries construction

We chose to perform the enrichment in pools of 48 individuals; as individuals were roughly organised by concentration, DNA amount in each pool was normalized to either 12ng/uL or to the concentration of the least concentrated individual of the pool, to maintain a good homogeneity in the representation of each individual in a pool. This approach allowed us to use the highest normalized concentration for most of the pools. Control individuals were included in each pool at a random location in normalized plates.

Allegro Targeted Genotyping v2 (TECAN, Redwood City, CA, USA) libraries were then built according to manufacturer protocol (REF du protocole, see Sung *et al*. 2024 for details). Briefly, normalized DNA of individuals was fragmented and end-repaired (Program 1, ran on an Eppendorf MasterCycler EP (Eppendorf AG, Hamburg, Germany), and individual barcodes ligated (Program 2). Hybridization was performed overnight (Program 3), with the final incubation step at 60° performed for more than 12h. Extension step was performed according to manufacturer.

Enriched, pooled libraries were then amplified to add the Metaplex barcode, added on a 48 individual pool basis, *i*.*e*. 4 pools of 48 individuals shared the same Metaplex barcode. Required PCR-cycles were determined by EvaGreen qPCR (MxPro, Agilent Technologies), and pools sharing the same number of determined PCR-cycles were amplified in the same PCR run (Program 4).

#### Libraries sequencing

Libraries were quantified by qPCR (KAPA Library Quantification Kit for Illumina Libraries - KapaBiosystems), and their profiles were assessed on the Agilent Bioanalyzer (Agilent Technologies, Santa Clara, CA, USA) using a High Sensitivity DNA kit. For a few libraries, profiles revealed the presence of residual primers/adapters; these librairies were re-purified as in the last step of the protocol. The libraries were then sequenced on an Illumina NovaSeq6000 instrument, on 2 lanes of a S4 flowcell (Illumina, San Diego, CA, USA), in PE150 mode. Produced reads were trimmed as described above for WGS sequencing.

#### Bioinformatics Data Analysis

Two paired-end fastq files were produced for each accession, with a total of 768 files. The fastq files were first checked for the quality of the raw reads using FastQC (Andrews, 2010), and low quality and short sequences were trimmed using Trimmomatic (Bolger et al. 2014) with the parameters LEADING:20 TRAILING:20 SLIDINGWINDOW:4:30 MINLEN:30. Reads were mapped to the Idéo genome assembly (Demenou et al., 2025) with BWA v0.7.15, then ProbeFilter (available at https://interval.bio/allegro_bioinformatics.html) was used to mask the region of each reads corresponding to the probe sequences, and duplicates marked using Picard MarkDuplicates. Variant calling was performed individually using GATK HaplotypeCaller (with options --min-pruning 3 --max-num-haplotypes-in-population 200 --emit-ref-confidence GVCF --max-alternate-alleles 1 -- contamination-fraction-to-filter 0.0 --native-pair-hmm-use-double-precision true --max-reads-per-alignment-start 0), then gvcf files were combined with Picard GatherVcf and genotypes called using GATK GenotypeGVCFs. Picard FilterVcf and home-made R scripts were then used to filter out SNP with DP<10; GQ<20. A custom-made Rscript was used to modify heterozygous genotypes with and AF < 0.2 or >0.8 to the corresponding homozygous.

#### Control samples repeatability

As Idéo individuals were included as control in the experiment, we calculated the repeatability of this experiment as a quality control of the SNP panel and Allegro technology. Eight genotyping reactions were performed (one in each of the height pools, see above), and compared. We used SNP with no missing data (29,219 SNP) and considered a SNP repeatable only if the height replicates has exactly the same genotype (0/0, 0/1 or 1/1).

#### Genetic diversity analysis

Markers and genotypes with more than 50% missing data were discarded. The remaining markers were imputed using Beagle v5.4 (Browning, 2008; Browning and Browning, 2016) with default parameters. The genotyping matrix was then filtered out for markers with minor allele frequency (MAF) less than 5%. To assess the density of markers in the genotyping tool, the distribution of the markers across the fifteen flax chromosomes was visualized. A Principal Component Analysis (PCA) was conducted on the matrix using the R package FactoMineR v2.11 (Lê et al., 2008) to visualize the diversity and clustering following the main flax type. For diversity parameters, the proportion of polymorphic SNP among all target SNP was computed; and the expected mean heterozygosity (Berg and Hamrick, 1997) for all the panel_384 were computed as:

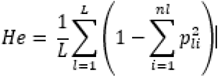

where L is the number of loci, n_l_ is the number of alleles at locus l, p_li_ is the relative frequency of the nbi-th allele at locus l.

## RESULTS AND DISCUSSION

### Sequencing, coverage and read counts statistics

In total, 1793 Gb of Illumina raw sequencing data was generated, representing 6 billion 2 x 150 bp paired-end sequences for the panel_96. An average of 126 million reads were generated per genotype, with the number of reads ranging from 73.4 million (for Linea_22) to 222.6 million (for ARETHA); representing a coverage of 25x to 76x (considering the Idéo genome size of 437 Mb). Reads counts, size and coverage are presented in Supplementary Table S4. After trimming with Trimmomatic, from 62 million (for Linea_22) to 189.6 million reads (for ARETHA) representing a coverage of 18x to 56x respectively, were kept. Average coverage before quality control (QC) and trimming was 42x (Standard Deviation SD = 4.21), dropping to 32x (SD = 3.25) and after QC and trimming. QC and trimming.

For the panel_77 accessions, 255 Gb Illumina raw sequencing data were downloaded from NCBI Sequence Read Archive. This represents an average 22 million of reads per genotype (number of reads per genotype ranging from 18k (for Viking) to 92 million (for CIli642), showing unbalanced data between genotypes. After trimming, from 18k (for Viking) to 24.6 million reads (for CIli1407) representing a coverage of 1x to 9x respectively, were kept. Average coverage before quality control (QC) and trimming was 7x (Standard Deviation SD = 2.47), dropping to 6x (SD = 1.74) and after QC and trimming (see Supplementary Table S5).

### De novo high-throughput SNP discovery and selection of 41k SNP

The goal is to develop a genotyping tool primarily for breeding European lines. Accordingly, we identified variants separately for the two panels. Of the raw reads for each accession (see Supplementary Table S4 and Table S5), more than 95.58% were mapped to the Idéo reference assembly for panel 77, compared to more than 97.67% for panel 96. SNP calling enabled the identification of 4.78 and 3.73 million biallelic SNPs in panel_96 and panel_77, respectively (see Supplementary Table S6). After filtering SNPs according to all criteria, 70,389 SNPs were retained for panel_96 and 3,946 for panel_77 (Table 1 and Supplementary Table S6). All 3,946 SNPs in panel_77 were retained for the probe design step (see Table 1). Of the 70,389 SNPs in panel_96, 18,829 in the genic region were retained and the remaining 18,746 were selected to ensure equivalent density along the chromosomes (see Table 1). A total of 41,521 SNPs were selected and sent to the Tecan company for probe design.

**Table 1.** The overall statistics for SNP selection and probe design success, and the distribution across the fifteen flax chromosomes.

| Chromosome ID | SNPs Selection |  |  |  |  | Probe design |  |
| --- | --- | --- | --- | --- | --- | --- | --- |
|  | Panel_96 | Panel_77 | TOTAL | Chromosome size | SNP density (SNP/bp) | Number of SNP COVERED | Number of SNP NOT COVERED |
| ALL | 37 575 | 3 946 | 41 521 | 437 Mbp | 11 000 | 35791 | 5 730 |
| Lus01_Ideo | 2176 | 236 | 2412 | 31 426 569 | 13 000 | 2 140 | 272 |
| Lus02_Ideo | 2538 | 464 | 3002 | 32 468 936 | 11 000 | 2 576 | 426 |
| Lus03_Ideo | 3370 | 748 | 4118 | 32 136 534 | 7 800 | 3 903 | 215 |
| Lus04_Ideo | 2264 | 210 | 2474 | 37 773 654 | 15 500 | 2 068 | 406 |
| Lus05_Ideo | 2332 | 200 | 2532 | 30 470 237 | 12 000 | 2 242 | 290 |
| Lus06_Ideo | 2811 | 84 | 2895 | 29 273 093 | 10 000 | 2 409 | 486 |
| Lus07_Ideo | 2789 | 333 | 3122 | 23 253 233 | 7 500 | 2 502 | 620 |
| Lus08_Ideo | 2021 | 102 | 2123 | 23 944 318 | 11 500 | 1 930 | 193 |
| Lus09_Ideo | 1502 | 107 | 1609 | 16 532 512 | 10 000 | 1 479 | 130 |
| Lus10_Ideo | 2466 | 246 | 2712 | 26 458 982 | 10 000 | 2 371 | 341 |
| Lus11_Ideo | 2375 | 105 | 2480 | 25 447 886 | 11 000 | 2 287 | 193 |
| Lus12_Ideo | 2865 | 535 | 3400 | 26 321 240 | 8 000 | 2 789 | 611 |
| Lus13_Ideo | 2316 | 121 | 2437 | 32 119 806 | 13 500 | 2 117 | 320 |
| Lus14_Ideo | 2819 | 74 | 2893 | 19 368 417 | 7 000 | 2 634 | 259 |
| Lus15_Ideo | 2744 | 118 | 2862 | 21 095 176 | 8 000 | 2 285 | 577 |
| Unknown CHR | 187 | 263 | 450 | 29 193 158 | 65 000 | 59 | 391 |

As shown in Figure 1, the selected SNPs were uniformly distributed across all chromosomes, except in the centromeric regions of many of them. Low density of SNP was also observed on one telomeric regions for chromosomes Lus5, Lus7 and Lus9.

**Figure 1.**
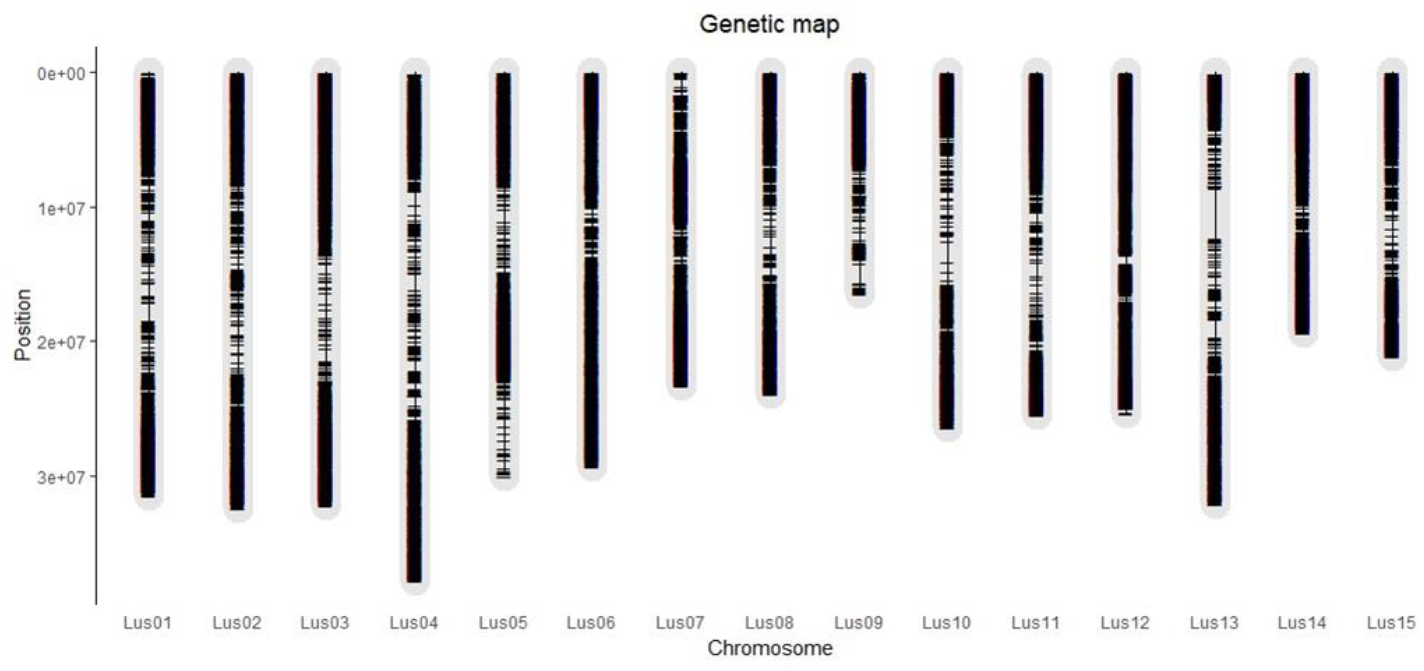
Physical position of the selected SNPs showing the density of the 41,521 SNPs along the fifteen flax chromosomes.

### Targeted SNP Custom probes design

Using the in-house pipeline, Tecan designed a couple of primers for each of the 41,521 target SNP taking into account context sequences and target SNP position. In total, 68,843 probes were designed, including 33,052 SNPs that were totally covered by two probes, 2,739 SNPs that were partially covered by one probe, and 5,730 SNPs that were not covered (Table 1). The success rate of probe design is 86% (35,791 SNPs). For the 5,730 SNPs for which no probe was designed, either no design was possible or none of the designs met all the requirements, including the BLAST hits being less than five.

### Validation of the genotyping tool

We tested the genotyping tool by genotyping all accessions of the panel_384. All accessions of this panel were sequenced for the target 35,791 SNPs in this genotyping experiment, following the Allegro targeted SNP genotyping protocol. The targeted sequencing coverage for each accession was approximately 100X. Figure 2 shows the distribution of the number of target SNPs based on the average depth without duplicate reads at the target SNP after read mapping. Therefore, after mapping the produced sequences against the Idéo genome assembly, the average SNP target coverage was found to be 68X once duplicate reads had been removed.

**Figure 2.**
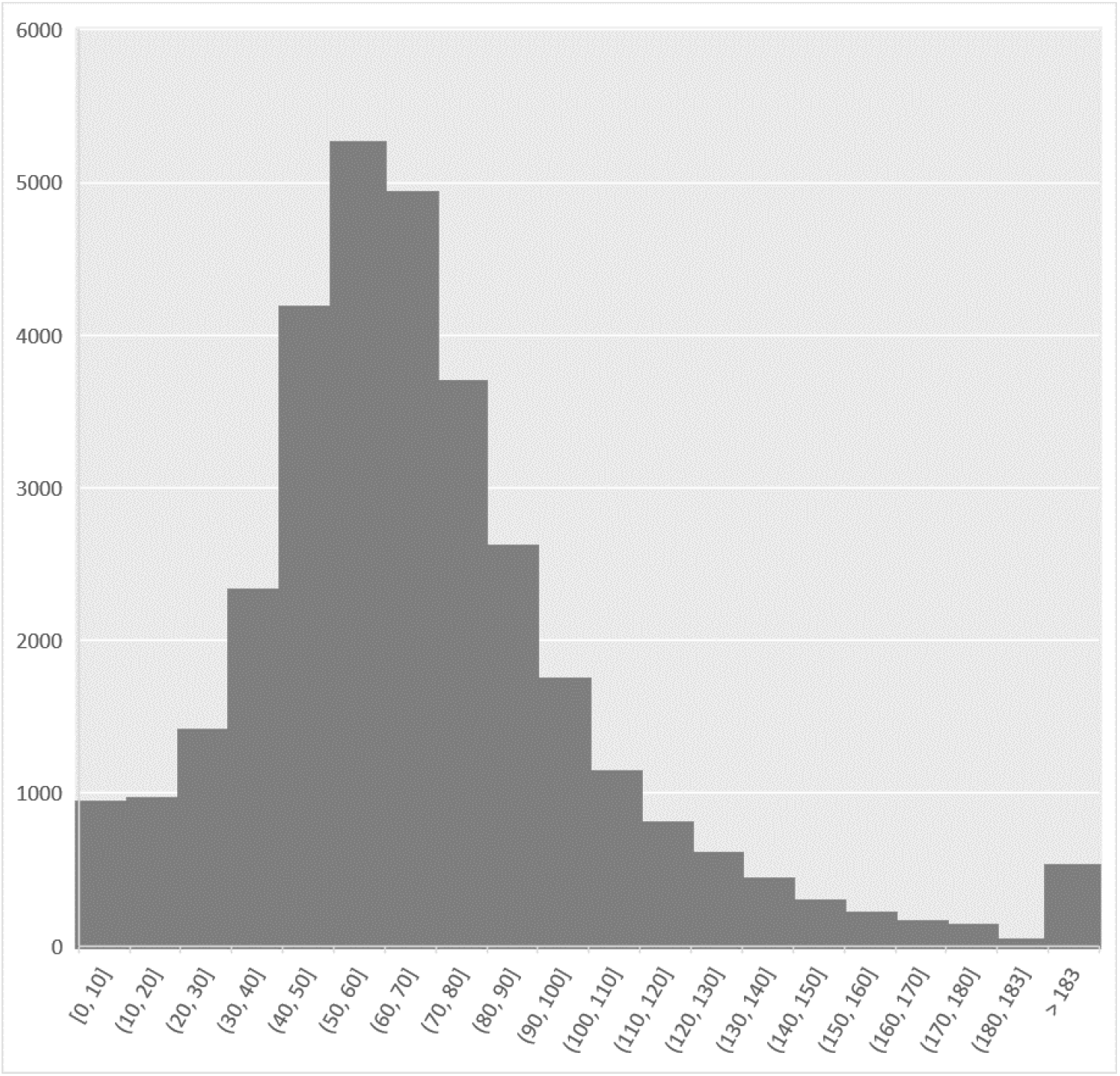
Distribution of number of target SNPs based on the average depth without duplicates reads at the target SNP.

As expected, all reads align with the Idéo genome, but some off-targets were observed. Off-target reads correspond to reads that map to a position in the genome other than the target SNP. For this reason, off-target reads were tagged, and only properly paired reads were used for genotype calling. Genotype calling was performed using the pipeline described above. As a result, of the 35,791 target SNPs included in the genotyping, 33,703 (94.16%) were variants, of which 33,175 (98.44%) were biallelic and 528 (1.56%) were triallelic (see Table 2). Of the target SNPs, only 2,088 (5.65%) failed to be genotyped, indicating a high success rate (see Table 2). At this step, 33,175 SNPs meet all conditions to be used on the genotyping tool.

**Table 2.** Summary of the number of target SNPs validated by the panel_384 genotyping.

|  | Quantity | Proportion of SNP (%) |
| --- | --- | --- |
| Nb of SNP Target | 35 791 | 100% |
| Nb of failed SNP target | 2 088 | 5.83% |
| Nb of Variant Target | 33 703 | 94.16% |
| Nb of Variant Target with more than one alternative allele | 528 | 1.56% |
| Nb of biallelic Variant Target | 33 175 | 98.44% |

When looking at the frequency parameters of the resulting genotyping matrix (33,175 SNPs for 376 accessions), we found that more than 25,000 SNPs have less than 10% missing data and that more than 22,000 SNPs have less than 20% heterozygosity. For the accessions, only eight individuals have more than 50% missing data (see Figure 3 and Supplementary Table S7). The genotypic frequency ranged from 1% to 82% for homozygotes for the reference allele (0/0), and from 0.2% to 26.3% for homozygotes for alternative allele. The percentage of heterozygotes ranged from 0.1% to 19.9% for the accessions (see Supplementary Table S7).

**Figure 3.**
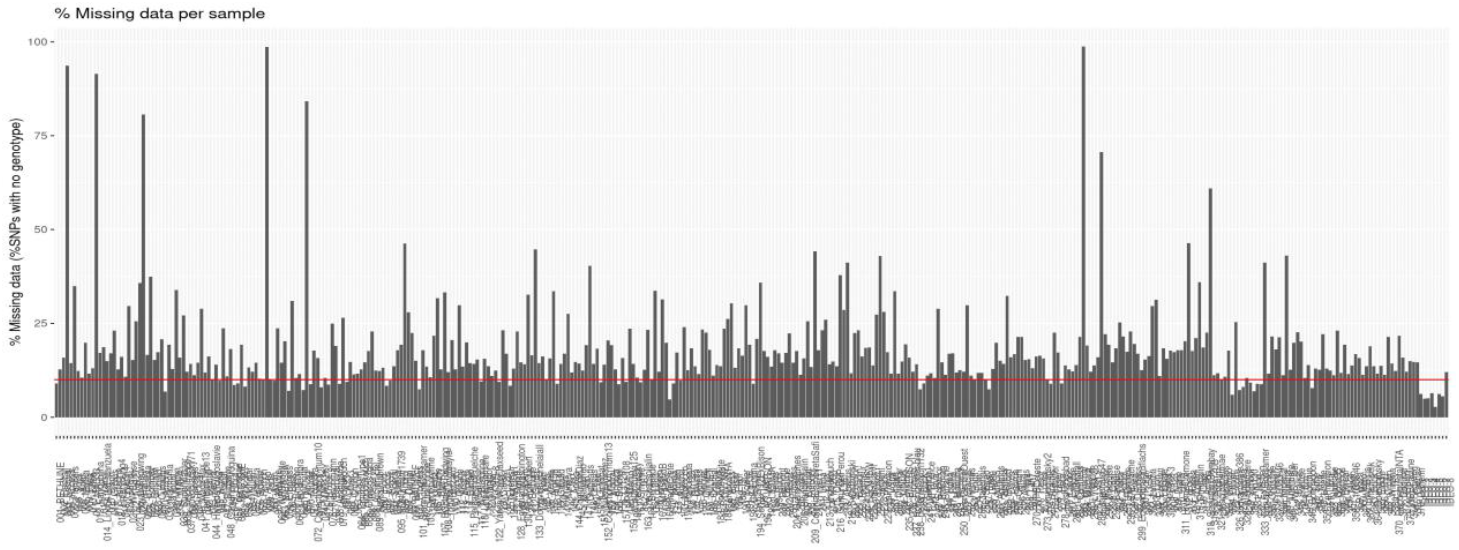
Percentages of missing data “samples” for biallelic target variants. The red horizontal bar indicates a missing data percentage of 10%.

### Repeatability of genotyping controls

In the genotyping experiment, the repeatability analysis of the genotyping tool was tested by genotyping eight replicates for the control Idéo. As result, 99.00% of the 29,219 SNPs with no missing data for all the eight replicates control have the same genotype and are repeatable for the Idéo control. However, it should be noted that there are 305 and 199 repeatable SNPs 0/1 and 1/1, respectively, between the Idéo controls.

### Genetic diversity analysis

The genotyping matrix obtained was filtered for missing data. Eight individuals with more than 50% missing data as well as 292 SNPs with more than 50% missing data, were removed before the matrix was imputed. The imputed matrix containing 33,175 SNP for 368 accessions, was filtered for minor allele frequency (MAF) less than 5% leading to a final matrix of 24,951 SNPs for 368 accessions. This final resulting matrix was used to compute the diversity parameters. Principal component analysis showing the distribution of the accessions in the validation panel based on the main flax type (Figure 4), demonstrates a strong genetic structure between oilseed and fiber types. It also enables us to distinguish dual-purpose types, which are located at the intersection of the two, and to classify unknown types into one of the remaining three categories. The average expected heterozygosity (He) computed for all 24,951 SNPs was 0.325.

**Figure 4.**
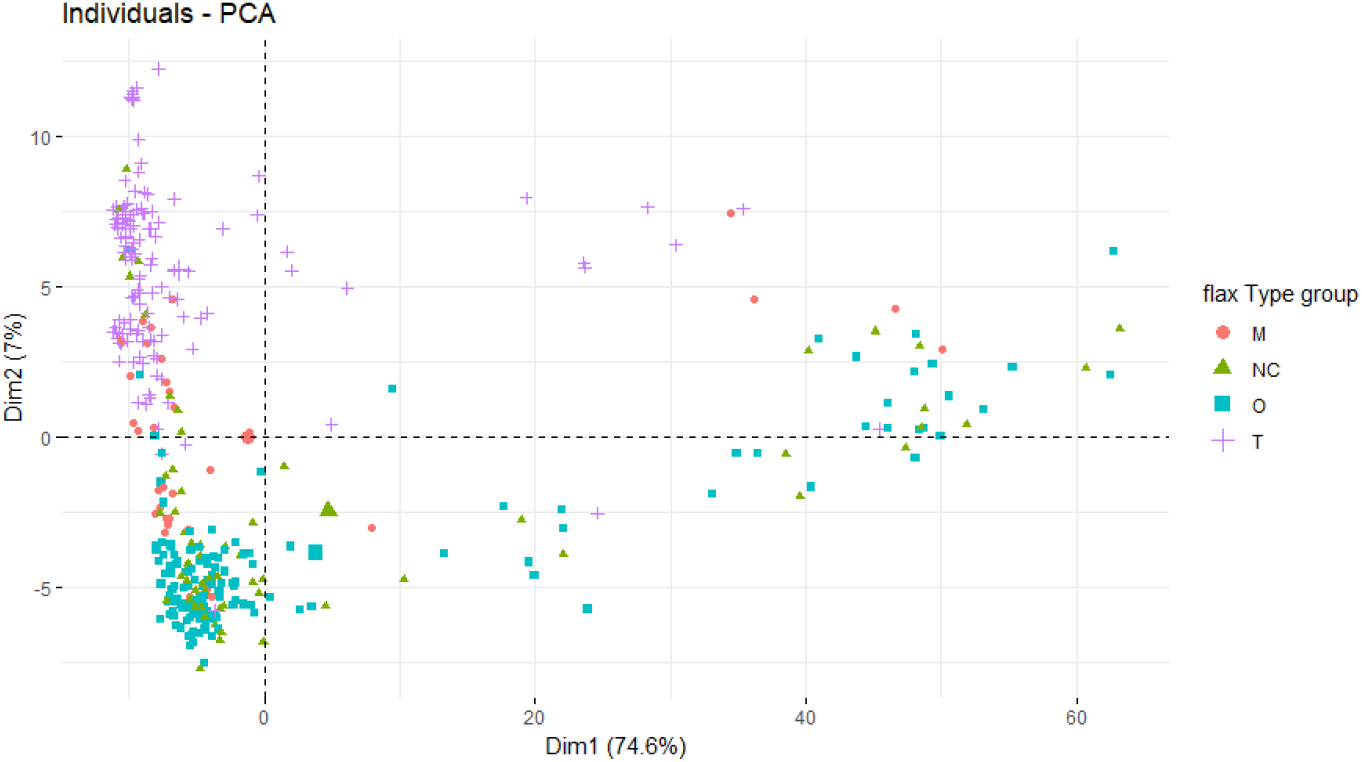
Principal component analysis (PCA) showing the distribution of the accessions in the validation panel based on the main flax type (M: dual-purpose; NC: unknown; O: oilseed; and T: fiber), on the first two axes.

## DISCUSSION

High-density SNP arrays have been widely used in economically important crop species enabling genotype characterisation, genetic studies, and the efficiency of plant breeding programmes. Even though European breeding companies have developed low- to medium-density in-house genotyping tools for cultivated flax, there are no high-density genotyping tools available to the general public. In this study, we used high-throughput SNP discovery methods to identify a set of SNPs representative of global diversity. We then selected more than 40,000 SNPs (up to 90% of the SNPs are characteristic of European diversity) to develop and validate a 30K target SNP genotyping tool for breeding and characterizing cultivated flax (Linum usitatissimum). We used Allegro targeted SNP technology for this.

The first step in this study was to select the panels to be used for variant calling. Panel 96 was selected primarily to represent the genetic diversity of European flax. Panel_77 was then selected to complement the first panel and add global diversity. Although the main goal was to develop genotyping for European flax breeding, we would also be able to analyse global diversity using the genotyping tool. Raw read sequences were generated for the panel_96 and were downloaded from the NBCI SRA database for the panel_77. We obtained an average sequencing coverage of 32x after trimming (range of 18x to 56x) for the panel_96; a value comparable to, or higher than, that reported in other studies. This is higher than the 15x obtained by Speck et al. (2022) and Guo et al. (2020) for flax, and the 18x used by Rimbert et al. (2018) for wheat. It is therefore highly suitable for variant identification analyses, allowing for robust results. Therefore, less data was available for the second panel_77 (less than 10x an average after trimming) compared to panel_96, and the variant calling parameters were adapted. SNPs calling was performed separately for the two panels, as goal is to develop a genotyping tool primarily for breeding European lines. This is because using both panels simultaneously would prioritise the identification of markers from worldwide diversity over those from European diversity. Of the raw reads for each accession (see Supplementary Table S4 and Table S5), more than 95.58% were mapped to the Idéo reference assembly for panel 77, compared to more than 97.67% for panel 96. This highlights a high mapping rate. Up to 4.78 million of candidate SNPs were identified for each panel (see Supplementary Table S6).

Using a stringent set of filtering criteria, 70,389 SNPs were retained for panel_96 and 3,946 for panel_77 (Table 1 and Supplementary Table S6).The low number of retained SNPs in Panel_77 is probably due to the lower sequencing coverage of its accessions compared to those in Panel_96 (see Supplementary Table S4 and Table S5). The majority of SNPs were lost during the SNP filtering step, with a significant drop occurring after filter 5 (Only SNPs with a percentage of heterozygosity below 20% and a homozygous genotype frequency between 5% and 95% are kept.) when the number of SNPs fell from 277k to just 9k (see Supplementary Table S6). In the end, 41,521 SNPs were selected. This represents an average density SNP of one SNP every 11 kb of the fifteen chromosomes, the chromosome Lus14 was the densest in SNPs with one SNP every 7 kb, while the chromosome Lus04 was the least dense (see Table 1 and Figure 1). Therefore, the selected SNPs were uniformly distributed across all chromosomes, except in the centromeric regions of many of them. Low density of SNP was also observed on one telomeric regions for chromosomes Lus5, Lus7 and Lus9. These results are probably due to either the chromosome regions mentioned above being repeats, and/or to low read sequence coverage during sequencing or after sequence mapping. It is well known that repeat regions experience either low or high coverage after reads mapping, as the same sequenced reads could map to many other positions within the repeats. Therefore, the depth after read mapping could fall outside the range of minimal and maximal depths. The set of selected SNPs was sent to the Tecan company for probe design. The success rate of probe design is 86% (35,791 SNPs), that is a good rate despite the level of genome duplication of flax (up to 73%, Demenou et al., 2025). For the next step of genotyping validation, the panel_384 was genotyped for the target 35,791 SNPs with successful probe design. A sequencing depth of 100x was expected following genotyping. However, mapping against the Idéo genome assembly yielded an average SNP target coverage of 68x once duplicate reads had been removed (see Supplementary Table S7). This is an acceptable average level of coverage for high-quality genotype calling, with a low level of missing data. Since all reads were mapped to the Idéo genome, some of the reads must have been mapped to positions other than the target positions. These off-target reads and the duplicate reads accounted for 326% of all reads produced. The genotyping matrix generated showed that for the panel_384, high-quality and discriminant markers were selected. A low average missing data of 18% was identified. Only eight accessions and 292 SNPs had more than 50% missing data. At this missing data level of 50%, It has been proven with our data that imputation can be performed with less than a 5% risk of introducing errors into the matrix. These results, showing 50% filtration, demonstrate the high-quality of the tool and the tool validation experiment, as very little data was lost due to missing data. Following the imputation of missing data, 24,951 SNPs have a minor allele frequency (MAF) greater than 5%, making them informative for genetic analyses and breeding. It is important to note that setting the MAF threshold at 1% increases the number of usable SNPs to 27,247. The frequency of heterozygote accessions for SNP marker ranged from 0 to 94.4%, with an average of 5.2%. This low overall heterozygosity rate was expected for homozygous lines, but it was high compared to that obtained by Rimbert et al. (2018) for wheat (1.5%). The difference must be explained by the presence of TRT accessions in panel_384 that were not considered as lines. The heterozygous genotypic frequency (the rate of SNP heterozygosity) then ranged from 0.3% to 19.9% for the accessions, which complies with the acceptable level of heterozygosity for genetic association analysis.

One of the main criteria for the success and quality of the developed genotyping tool is repeatability. In the genotyping validation experiment, 99.00% of SNPs were repeatable for the eight Idéo replicates controls. This is satisfactory results about the use of the genotyping tools at large-scale on larger panels. Finally, the generated genotyping data enabled the computation of diversity parameters, demonstrating the strength of this matrix in classifying the accessions in this panel, and furthermore the strength of the target SNPs genotyping tool developed for further genetic analysis.

## CONCLUSION

This study developed and validated the potential application of the Allegro targeted SNP genotyping tool in genomic research, genotype characterization and genetic improvement of the European cultivated flax. The tool contains 35,791 SNPs, covering the fifteen chromosomes and the entire genome, and demonstrates high polymorphism and excellent genotyping accuracy. Of this list of SNP, 24,951 high-quality SNPs were validated on the panel_384. This tool exhibit also a rate of 99% of repeatability, a good quality criterion for the use at large scale. It would provide a strong support for genomic and genetic research and would be useful for the European breeding companies for genetic improvement, germplasm resource management.

## ACKNOWLEDGMENTS

The authors wish to thank, Yann Flodrops for his help conceiving the GenoFLAX projet (sequencing, assembly and annotation of European flax cultivars and development of genotyping tool), Aurélie Canaguier (EPGV) for her help in providing the sequencing data, the CEA-IbFj/Genoscope sequencing platform staff for their help and Henri Desaint for constructive discussions about the selection of target SNPs. They are grateful to Isabelle Chaillet, the flax germplasm collection manager at Arvalis Institute, for her help in selecting panel 384.

## AUTHOR CONTRIBUTIONS

B.D. conceived the research. C.P.P. selected the panel_96, B.D. selected the panel_77; C.P.P. and B.D. selected the panel_384; C.P.P. provided the plant material (seeds and DNA extract) for the panel_96 and 384. D.D.H. and P.F.R. supervised the probe design; Tecan designed the probes and performed the synthesis of *in solution* probes, D.D.H., P.F.R. and B.D. validated the design. D.D.H. and P.F.R. supervised the wet labo manipulations and provided Illumina sequences data for the panel_96 and the genotyping data for the panel_384. B.D., D.D.H. and P.F.R. analysed the data. B.D. prepared all figures; B.D., C.P.P., P.F.R. and D.D.H. wrote the manuscript. All authors read, edited and contributed to the manuscript.

## CORRESPONDING AUTHOR

Correspondence to Boris B. Demenou (*)*

## SUPPLEMENTARY MATERIALS

**Table S1**. List of 96 contemporary European breeding accessions, representing the diversity panel for SNP identification

**Table S2**. List of 77 accessions retrieved from the collection of Guo et al. (2020), representing the second panel of SNP identification

**Table S3**. List of 376 accessions representing the validation panel

**Table S4**. The overall statistics of reads data generated from WGS re-sequenced for the 96 accessions of panel 96

**Table S5**. The overall statistics of reads data downloaded from SRA database, for the 77 accessions of panel_77

**Table S6**. The number of remaining SNPs for each filtering step

**Table S7**. Genotypic frequency of accessions in panel 384, computed from data on 33,175 SNPs

